# Thioridazine as an Anticancer Therapeutic: Interplay with the Isoprenoid Biosynthetic Pathway

**DOI:** 10.1101/532788

**Authors:** Jillian S. Weissenrieder, Jessie L. Reed, Jeffrey D. Neighbors, Raymond J. Hohl

**Author notes:** Corresponding author: Raymond J. Hohl, M.D., Ph.D. Penn State Cancer Institute Penn State College of Medicine 500 University Drive, Mail Code CH72 Hershey PA 17033-0850.

## Abstract

Glioblastoma multiforme is a form of cancer with poor survival prognosis and few treatment options. The cerebrovascular barrier complicates the delivery of chemotherapeutic agents and contributes to poor treatment response in patients with this disease. Recently, dopamine D_2_ receptor antagonizing compounds, including the FDA-approved phenothiazine, thioridazine, were identified as potential anticancer therapeutics, but their mechanism of action is as yet poorly understood. We investigated the hypothesis that the cytotoxicity of thioridazine may be tied to disruption of lipid metabolism, specifically the synthesis of isoprenoids and cholesterol by the isoprenoid biosynthetic pathway. We show that, while pathway inhibitors lovastatin and zoledronate can sensitize U87MG and U251MG cells to thioridazine treatment, the addition of pathway intermediates cannot prevent thioridazine’s cytotoxic effects. Treatment with methyl-schweinfurthin G, which is known to disrupt lipid trafficking, is able to sensitize these cell lines as well, suggesting that cholesterol availability or localization may be involved in these effects. However, all measured effects were of very small, biologically insignificant magnitude and thus findings are of limited utility.

## Introduction

Glioblastoma multiforme (GBM) is one of the most common central nervous system malignancies (Brodbelt et al., 2015; Alexander and Cloughesy, 2017; Ostrom et al., 2018). While metastasis of GBM is rare, survival rates are dismal, with the vast majority of patients surviving under two years after diagnosis (Koshy et al., 2012; Brodbelt et al., 2015; Ostrom et al., 2018). The current standard of care, consisting of maximal surgical resection, radiotherapy, and chemotherapy with temozolomide (Alexander and Cloughesy, 2017). While many types of cancer have benefited strongly from the development of targeted therapeutics, many of these have shown limited efficacy in GBM due to the exclusion of therapeutics by the blood-brain barrier (BBB) and the immune privilege of the environment (Maxwell et al., 2017; Touat et al., 2017). Therapy is also complicated by inter- and intra-patient tumor heterogeneity (Sottoriva et al., 2013; Patel et al., 2014; Morokoff et al., 2015) and diffuse tumor margins (Chaichana et al., 2014; Pessina et al., 2017). A novel treatment method using alternating electric fields has shown some efficacy, but no survival improvement over the current standard of care (Stupp et al., 2012). To improve patient outcomes, new treatment options are needed.

Lipid modulation shows promise as an anti-GBM therapeutic target. Glial cancers like GBM are known to be highly dependent on lipid metabolism, cholesterol levels, and liver X receptor (LXR) function, suggesting lipid-targeting therapeutics, especially those which modulate cholesterol synthesis or trafficking, could have potential in the treatment of GBM (Villa et al., 2016). Cholesterol is a product of the isoprenoid (or mevalonate) biosynthetic pathway (Figure 1), which may be pharmacologically inhibited by statins, such as lovastatin. Statins inhibit the rate limiting step of cholesterol synthesis catalyzed by HMGCoA reductase. Other enzymes within this pathway may also be modulated with chemical inhibitors, allowing the role of individual pathway intermediates in a given phenotype to be readily discerned by experimentation.

**Figure 1.**
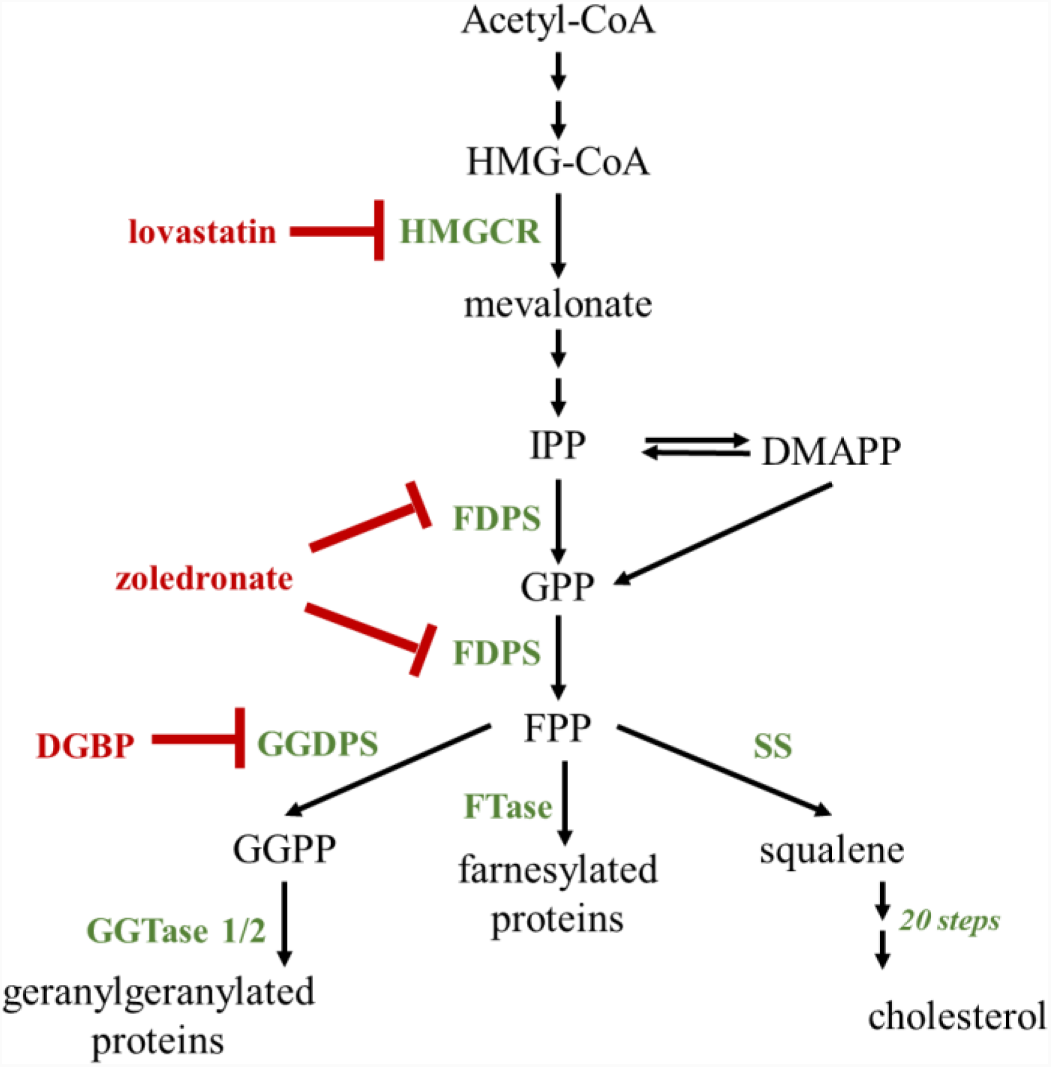
The isoprenoid pathway is involved in the synthesis of cholesterol and isoprenyl groups. It may be inhibited at key steps. Notably, the statins, such as lovastatin, inhibit the rate limiting step near the beginning of this pathway, synthesis of mevalonate from HMG-CoA via HMG-CoA reductase (HMGCR). Zoledronate inhibits the conversion of isopentenyl pyrophosphate (IPP) to geranyl pyrophosphate (GPP) and farnesyl pyrophosphate (FPP) by farnesyl diphosphate synthase (FDPS). The pathway branches out at FPP, which can be used to isoprenylate proteins through the action of farnesyl transferases (FTases), as a substrate to synthesize geranylgeranyl pyrophosphate (GGPP), or as a substrate for the committed step of cholesterol synthesis by squalene synthase (SS). Digeranyl bisphosphonate (DGBP) inhibits the conversion of FPP to GGPP by geranylgeranyl diphosphate synthase (GGDPS), reducing substrate availability for geranylgeranyl transferases 1 and 2 (GGTase 1/2).

Recently, dopamine D_2_receptor (D_2_R) antagonists have been identified as potential anticancer therapeutics (Rho et al., 2011; Sachlos et al., 2012; Yeh et al., 2012; Cheng et al., 2015; Huang et al., 2016). In the case of GBM, their ability to pass through the blood-brain barrier makes them particularly attractive compounds (Batash et al., 2017; Touat et al., 2017; Harder et al., 2018). Numerous compounds of this class are approved by the Food and Drug Administration for treatment of mental illnesses, such as schizophrenia. A number of these D_2_R antagonists, especially those of the phenothiazine chemotype, have been shown to induce apoptosis (Gil-Ad et al., 2004), decrease survival signaling (Park et al., 2014), and reduce tumor size in xenograft models (Shin et al., 2013). All of these effects are desirable in an anticancer therapeutic.

Interestingly, D_2_R antagonists are also known to disrupt lipid synthesis and trafficking (Masson et al., 1992). It is not known if this dysregulation is required for their anticancer activity. By the early 1990s, it was evident that cytotoxic concentrations of the phenothiazine D_2_R antagonist, chlorpromazine, inhibited sphingomyelinase activity and induced accumulation of unesterified cholesterol in large droplets within the cell (Masson et al., 1992). Treatment with D_2_R antagonists such as haloperidol and clozapine increased sterol responsive element binding protein 1/2 (SREBP) protein levels and induced upregulation of sterol-responsive genes such as *HMGCR, APOE, ABCA1, LXR1/2*, and others in the context of GaMg glioma cells (Ferno et al., 2006; Vik-Mo et al., 2009). This increase in mRNA may lead to a buildup of synthetic pathway intermediates, but appears to block cholesterol synthesis (Sanchez-Wandelmer et al., 2010; Canfran-Duque et al., 2013).

Here, we more closely describe the role of the isoprenoid pathway in the mechanism of action for the phenothiazine D_2_R antagonist, thioridazine. Through a series of experiments involving supplementation with isoprenoid pathway intermediates and cotreatment with isoprenoid pathway inhibitors, we show that the anticancer efficacy of thioridazine is not reliant upon isoprenoid pathway disruption.

## Methods

### Cell culture and reagents

Cell lines U87MG and U251MG were purchased from American Type Culture Collection (ATCC, Manassas, VA). Both lines were cultured in MEM (Gibco, Waltham, MA) plus 10% FBS (HyClone, Logan, Utah) at 37°C and 5% CO_2_. Cell lines were used within 20 passages of receipt.

Thioridazine (Sandoz Pharmaceuticals, Holzkirchen, Germany) was maintained as a 10 mM stock in dimethyl sulfoxide at −20°C. Farnesol, geranylgeraniol, and mevalonate (1M stock) were obtained from Sigma Aldrich (St. Louis, MO). Lovastatin was activated as previously described (Dong et al., 2009). DGBP was synthesized as previously described (Shull et al., 2006) and dissolved in water. Likewise, methyl schweinfurthin G was synthesized (Koubek et al., 2018) and protected from light at all times. All compound stocks were maintained at −20°C at 10 mM in DMSO unless otherwise noted.

### MTT assay

MTT assays were carried out as previously described (Weissenrieder et al., 2018). Briefly, U87MG and U251MG cells were plated at 4000 cells/well in 96 well plates and allowed to seat overnight before treatment with 100 µL treatment media per well. Treatments were mixed in phenol red free MEM (Gibco, Waltham, MA) + 10% FBS. After 44 h of incubation at 37°C and 5% CO_2_, 10 µL of 5 mg/mL MTT (3-(4,5-dimethylthiazol-2-yl)-2,5-diphenyltetrazolium bromide, Life Technologies, Waltham, MA) was added to each well. At 48 h, 100 µL of stop solution (10% 1 N HCl, 10% Triton X–100, 80% 2-propanol) was added and incubated at 37°C overnight, then A570 (test) and A690 (reference) were read on a Spectramax i3x (Molecular Devices, Sunnyvale, CA). Absorbances were normalized to vehicle controls.

### Statistics

All data is representative of 2-3 independent experiments in triplicate. Concentration response curves were generated via non-linear regression and compared with GraphPad Prism 8 (San Diego, CA). Flow cytometric data was analyzed via chi-square.

## Results

### Cotreatment with isoprenoid pathway inhibitor lovastatin sensitizes GBM cells to thioridazine treatment

To observe if the cytotoxic effects of thioridazine were due to isoprenoid pathway alterations, we used 48 h MTT assays under concomitant treatment with pharmacological inhibitors of the pathway. For these experiments, we used 30 µM digeranyl bisphosphonate (DGBP), 10 µM zoledronate, and 3 µM activated lovastatin. At these concentrations, these compounds are known to selectively inhibit stages of the isoprenoid pathway (Figure 1).

Lovastatin is a commonly used competitive inhibitor of 3-hydroxy-3-methylglutaryl-CoA reductase (HMGCR), the rate-limiting enzyme of the isoprenoid pathway. Treatment with lovastatin reduces substrate synthesis and availability for all future pathway enzymes, decreasing both cholesterol synthesis and isoprenoid pyrophosphate availability. Zoledronate, a bisphosphonate, acts downstream of lovastatin to reduce farnesyl pyrophosphate (FPP) synthesis and modulate downstream enzymes, such as farnesyl transferases and squalene synthase (Dunford et al., 2001). This represents a branch point in the isoprenoid pathway, where FPP may be used to isoprenylate proteins, synthesize geranylgeranyl pyrophosphate (GGPP), or begin the committed cholesterol biosynthetic process. Farnesyl transferases require the farnesyl pyrophosphate (FPP) substrate generated by the isoprenoid pathway to post-translationally modify, and thus regulate, small GTPases. These proteins, including Ras family members, are known to contribute to cancer development and metastasis. Squalene synthase, however, requires FPP for the first committed, rate-limiting step to synthesize cholesterol, itself a required component of cell membranes. Downstream of FPP synthesis, DGBP selectively inhibits the synthesis of GGPP by geranylgeranyl diphosphate synthase (GGDPS), reducing the substrate availability for the isoprenylation of small GTPases in the Rho and Rab families by their respective transferases (geranylgeranyl transferase 1 and 2, respectively) (Wiemer et al., 2007; Wiemer et al., 2011). This can disrupt cytoskeletal regulation, thus potentially contributing to dysregulation of metabolic processes and trafficking of required metabolic and structural substrates. These enzymes are later-stage enzymes in the isoprenoid pathway, subject to the regulation of upstream enzymes such as 3-hydroxy-3-methylglutaryl-CoA reductase (HMGCR).

Metabolic activity of both U87MG and U251MG GBM cell lines were examined with 48 h MTT assays. We observed a small, yet statistically significant, reduction in inhibitory concentration (IC_50_) for thioridazine for cells cotreated with lovastatin and zoledronate (p<0.0001), but not for cells cotreated with DGBP (p>0.05) (Figure 2A-B). This suggests that the isoprenoid biosynthetic pathway may be involved in the mechanism of action for thioridazine, since an inhibitor of this pathway is able to sensitize cells to thioridazine treatment. However, the disruption of small GTPase regulation by geranylgeranylation may not be involved in this process, indicating that the earlier steps in the biosynthetic pathway may be more critical for cell survival under treatment with thioridazine.

**Figure 2.**
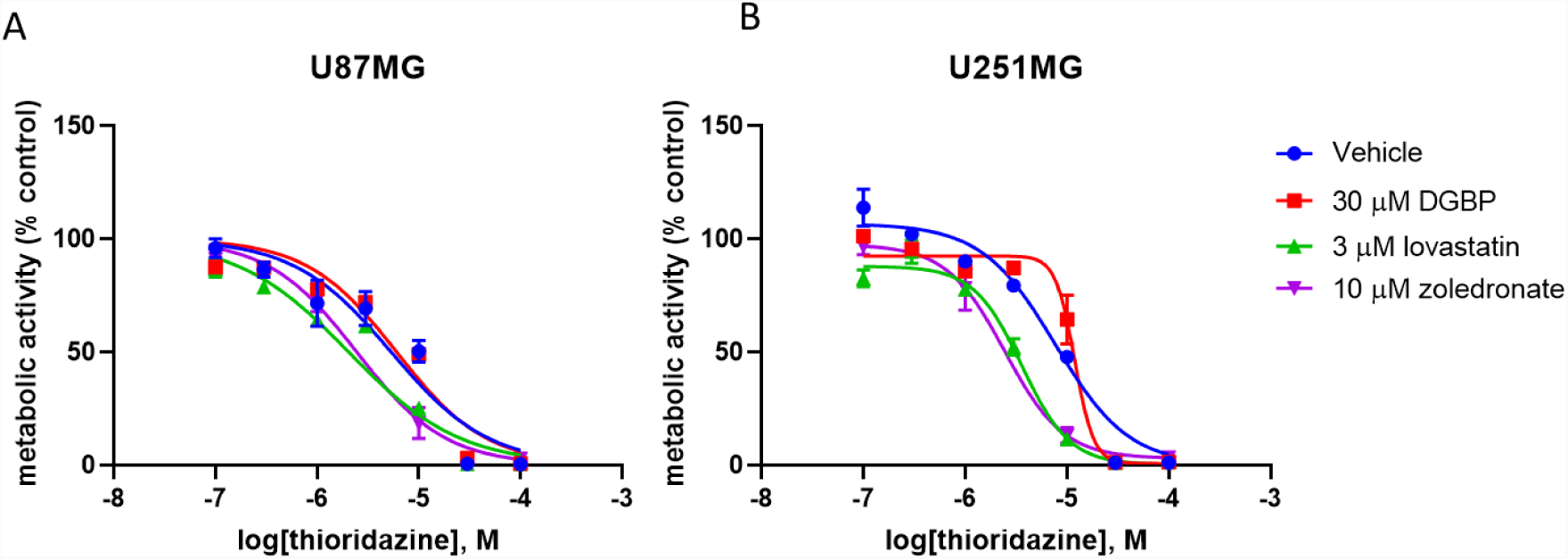
Isoprenoid pathway inhibitors sensitize GBM cells to thioridazine. In a 48 h MTT assay, 3 µM lovastatin and 10 µM zoledronate sensitize U87MG (A) and U251MG (B) cells to thioridazine treatment (p<0.0001). However, 30 µM digeranyl bisphosphonate has no significant effect on the sensitivity of these cell lines.

### Exogenous supplementation of isoprenoid pathway intermediates does not protect GBM cells from thioridazine treatment

Having observed that isoprenoid biosynthetic pathway inhibition could sensitize GBM cell lines to thioridazine treatment, we then examined whether the addition of pathway intermediates was sufficient to prevent the cytotoxic effects of thioridazine. This process is helpful to control for any non-selective effects of the pharmacological inhibitors in Figure 2. For FPP and GGPP, we supplied the alcohols of these pyrophosphates, which are more chemically stable and membrane permeable than the highly charged pyophosphates. When we treated cells with thioridazine and 5 mM mevalonate, 30 µM farnesol, or 20 µM geranylgeraniol, we saw no significant increases in IC_50_(Figure 3A-B). Indeed, supplementation with pathway intermediates farnesol and geranylgeraniol actually increased cytotoxicity (p<0.0001 for both). This may indicate that supplementation induced feedback inhibition on the isoprenoid pathway or that the protective effects of lovastatin and zoledronate are due to off-target interactions. However, the effects of mevalonate were more complex. Addback of mevalonate did not significantly sensitize U87MG cells to thioridazine, and it had a very slight, but statistically significant, protective effect on U251MG. This difference may be due to metabolic differences in the cell lines, but complicates findings.

**Figure 3.**
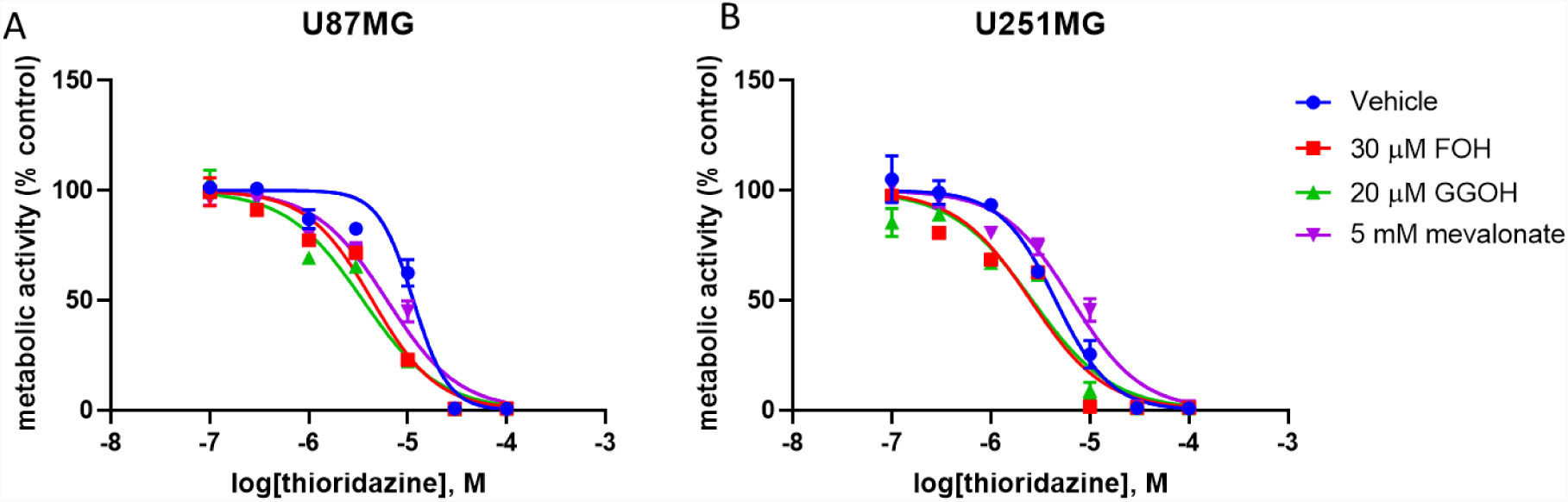
Exogenous supplementation of isoprenoid pathway intermediates does not have a pronounced effect on GBM cell sensitivity to thioridazine. Addback of geranylgeranyl and farnesyl moieties induced a low-magnitude, but highly significant (p<0.0001) sensitization of U87MG (A) and U251MG (B) cells to thioridazine treatment, according to 48 h MTT data. 5 mM mevalonate had no effect on U87MG, but was slightly protective for U251MG (p<0.05).

### Cotreatment with methyl-schweinfurthin G sensitizes cells to thioridazine treatment

We also ascertained the effects of cotreatment with a novel anticancer therapeutic from the schweinfurthin family, methyl-schweinfurthin G (MeSG). These compounds are known to have potent anticancer effects in the sub-micromolar range in models of central nervous system malignancies such as GBM (Beutler et al., 1998). While their anticancer mechanism of action is currently unknown, this class of compounds is known to disrupt lipid trafficking and metabolism in cell culture systems (Koubek et al., 2018). When we cotreated U87MG and U251MG cells with 100 nM MeSG, we saw a half-log order shift suggesting a more robust sensitization of these cell lines to thioridazine (Figure 4A-B). This suggests that perhaps the localization and/or concentration of cholesterol, its metabolites, or its precursors may affect sensitivity of cell lines to thioridazine treatment.

**Figure 4.**
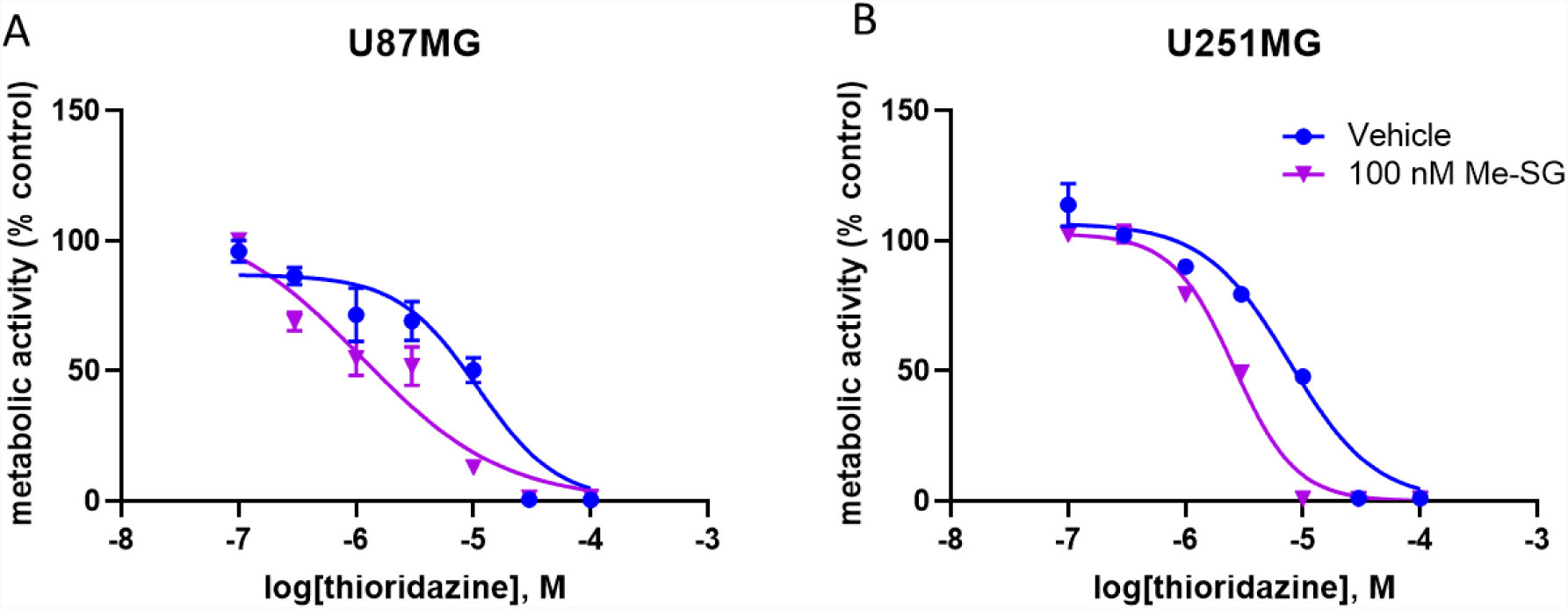
Cotreatment with 100 nM methyl-schweinfurthin G sensitizes GBM cell lines to thioridazine. In a 48 h MTT, U87MG (A) and U251MG (B) cell lines were significantly sensitized to thioridazine by 100 nM concentrations of the anticancer candidate compound, methyl-schweinfurthin G, which induced a half log order increase in sensitivity (p<0.0001).

## Conclusions

Taken together, our results indicate that depletion of isoprenoid pathway intermediates via upstream inhibition may increase sensitivity of the GBM cell lines, U87MG and U251MG, to the anticancer activity of thioridazine. However, treatment with these intermediates added back to the media is not protective. This suggests that, while this pathway may be perturbed to sensitize cells, its products are not sufficient to protect GBM cells from the cytotoxic effects of thioridazine. Such findings may be rationalized in a number of ways. For one, a fully functional isoprenoid pathway may provide some protection against thioridazine, but low level function without the addition of pathway intermediates may be sufficient for this effect. In this paradigm, only profound, near-total blockade of the pathway is sufficient to have an effect. Another option is that such effects are dependent upon cholesterol concentration, and cholesterol is not synthesized quickly enough from intermediates to protect against cotreatment with thioridazine, which may have cytotoxic effects at much earlier time points. If so, addbacks could cause feedback inhibition which would effectively reduce pathway flux while thioridazine is inducing cytotoxicity.

Future work may further clarify the role of the isoprenoid biosynthetic pathway in thioridazine response. This work may include quantitation of cholesterol and isoprenoid pathway intermediates under treatment with thioridazine or related compounds. It may also involve carrying out experiments with lipid depleted medium, cholesterol extraction with methyl-β-cyclodextran, or cholesterol supplementation to more closely observe the role of the cholesterol synthesis branch of this pathway. It may also be instructive to observe the effects of isoprenoid pathway modulation and thioridazine treatment in the context of cancer stem cell enriched populations, as may be modeled by spheroid cultures. These cultures appear to better model patient responses to compounds and may provide a clearer picture of the *in vivo* effects of these treatments (Lee et al., 2006).

Importantly, it is imperative to note that while the effects of isoprenoid biosynthetic pathway inhibitors are statistically significant, the magnitude of the effect is negligible in all experiments except perhaps for the experiments involving MeSG, where the curves are separated by half a log order. These findings may thus assist researchers in understanding the cytotoxic cascade initiated by thioridazine, but they likely have no direct translational applicability.

Current findings support a conclusion that thioridazine induces a metabolic crisis which may relate to lipid metabolism. However, these effects are not causative of cytotoxicity, but rather an effect of it.

## Acknowledgements

We would like to thank Dr. Richard B. Mailman for his pharmacological insight on this project.

## Respective Contributions

JSW carried out all experiments with the assistance of JLR. JSW wrote the manuscript. JDN and RJH provided experimental guidance and edited the manuscript.

